# Morphine alleviates pain after heart cryoinjury in zebrafish without impeding regeneration

**DOI:** 10.1101/2020.10.01.322560

**Authors:** Sara Lelek, Mariana Guedes Simões, Bo Hu, Ahmed M.A. Alameldeen, Maciej T. Czajkowski, Alexander M. Meyer, Fabienne Ferrara, Jan Philipp Junker, Daniela Panáková

## Abstract

Nociceptive response belongs to a basic animal behavior facilitating adaptability and survival upon external or internal stimuli. Fish, similarly to higher vertebrates, also possess nociceptive machinery. Current protocols involving procedures performed on adult zebrafish including heart cryoinjury do not, however, take into account the adverse effects including pain that may potentially arise from these methodologies. Here, we assess the effect of two analgesics, lidocaine and morphine, followed after heart cryoinjury in zebrafish. Monitoring swimming behavior together with histology and gene expression analysis at the single cell level using scRNA sequencing and RNAscope fluorescent *in situ* hybridization technology, we show morphine, but not lidocaine, significantly improves animal welfare 6 hours post-cryoinjury, without impairing the heart regeneration process. Altogether, we propose morphine to be considered as the analgesic of choice to reduce post-surgical pain in adult zebrafish.

**Highlights:**

- Cryoinury could be considered as a potential noxious stimulus in adult zebrafish.
- Morphine but not lidocaine treatment effectively alleviates noxious effects post-cryoinjury.
- Lidocaine treatment delays heart repair and regeneration.
- 6 hours Morphine treatment after cryoinjury does not impede heart regeneration.

## Introduction

A mounting body of evidence suggests fish including zebrafish react to harmful stimuli by changing their behavior. This is manifested by the significant reduction in the normal activity and the swimming frequency as well as the increase in the ventilation rate (Reilly et al., 2008). Despite being frequently used as an animal model, studies elucidating the pain and pain relief in zebrafish are scarce (reviewed in Chatigny et al., 2018). Most of the current methodologies and protocols using the adult zebrafish do not take into account the possible animal discomfort associated with pain. Therefore, there is a great demand, both in ethical and legal sense, to revise commonly used protocols and improve the standards of animal welfare for the adult zebrafish model.

To date only few systematic studies describing the effect of analgesics and their efficacy to relieve discomfort and pain in fish are available (Chatigny et al., 2018; T. Martins et al., 2016). The most investigated pain-relieving drugs thus far have been local anesthetics, NSAID (non-steroidal anti-inflammatory drugs), and opioids in two fish species: rainbow trout and zebrafish (Chatigny et al., 2018). Local anesthetics interrupt nerve conduction by inhibiting the influx of sodium ions through voltage-gated sodium channels in axonal membranes (Schwarz et al., 1977). Fish are commonly treated with the local anesthetic tricaine methanesulfonate (or MS222) to induce general anesthesia (Sneddon, 2012), which is not sufficient to manage adverse effects associated with stress and/or pain. Lidocaine is another local anesthetic tested as a systemic pain reliever in humans, rodents and fish (Chatigny et al., 2018; Mao and Chen, 2000; Schwarz et al., 1977). Although lidocaine may ameliorate stress- and/or pain-related behavioral changes in zebrafish adults after certain procedures (Deakin et al., 2019), its adverse effects have been also recently reported in embryos as well as in adults. For instance, lidocaine was shown to exacerbate the symptoms associated with the bipolar disorder in embryos (Ellis and Soanes, 2012), and to induce more sedative-like effects in embryos (Lopez-Luna et al., 2017) as well as in adults through inhibition of acetylcholine activity (de Abreu et al., 2018).

Opioids are the most common analgesics used in fish. Their mode of action is to interact with µ, δ, or κ opioid receptors, where they mimic the actions of endogenous opioid peptides (Chatigny et al., 2018). Opioids increase K^+^ efflux or reduce Ca^2+^ influx, resulting in impeding nociceptive neurotransmitter release. Like lidocaine, morphine treatment seems to also induce contrasting effects (Chatigny et al., 2018; Lopez-Luna et al., 2017). On one hand, a low dosage of morphine led to hyperactivity in zebrafish embryos, while a high dosage reduced their swimming activity (Lopez-Luna et al., 2017). In adults, morphine alleviated noxious effects at high dosages (Deakin et al., 2019). Additionally, the morphine treatment may cause a number of potential side effects, including those associated with the cardiovascular and respiratory system (Chatigny et al., 2018). The inconsistences in the effects of the analgesics may be among others a consequence of differences in concentrations used or time of the treatment creating a great demand for systematic testing of analgesics after noxious stimuli.

The adult zebrafish has become a popular animal model to study vertebrate regeneration, including heart regeneration. Current methodologies inducing the injury of the heart tissue include resection, cryoinjury, genetic ablation and hypoxia/reoxygenation (González-Rosa et al., 2017; Marques et al., 2019). The cryoprobe injury model is based on rapid freezing-and-thawing of the heart tissue, resulting in the cell death of about 20% of cardiomyocytes (CM) of the ventricular wall (Figure 1 A, B) (Chablais and Jazwinska, 2012; Chablais et al., 2011; González-Rosa and Mercader, 2012). This procedure causes local damage to all cardiac cell types and leads to a transient fibrotic tissue deposition reminiscent of the fibrotic scar formed in mammals after myocardial infarction (Kikuchi, 2014). In general, within 30 to 60 days after injury the heart is fully regenerated (González-Rosa et al., 2017; Marques et al., 2019). Although the cryoinjury method has been widely used to induce cardiac injury in sedated zebrafish using tricaine methanesulfonate, analgesics were never tested during this procedure. Moreover, their effect on fish welfare after cryoinjury and on the regenerative process is unknown.

**Figure 1.**
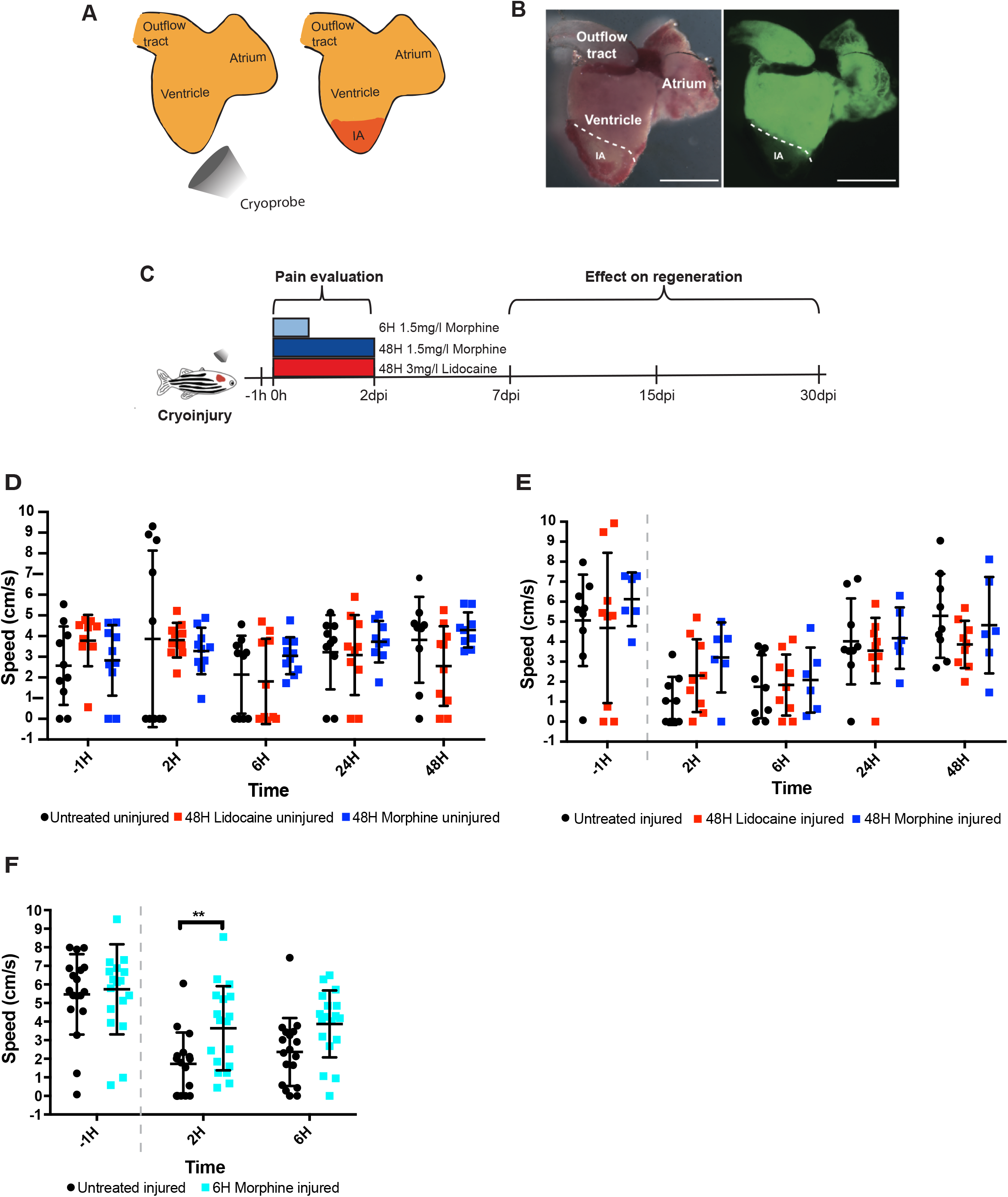
Morphine treatment alleviates pain after heart cryoinjury. (**A, B**) Cryoinjury induces the damage of about 20% of tissue in the heart ventricle. Schematic representation of the injury procedure (**A**), and injured *Tg(myl7:EGFP)* heart (**B**) 3 days post injury (dpi). Scale bar, 500 μm. (**C**) Experimental design of the study. (**D**) Effect of analgesic treatment on the swimming behavior of the uninjured fish. Swimming speed was assessed in fish one hour prior to analgesic treatments (−1H), and at 2, 6, 24, and 48H of treatment in untreated control fish (n=10, black circles), lidocaine-treated fish (n=10, red squares), and morphine-treated fish (n=10, dark blue squares). (**E**) Effect of the analgesic treatment on the swimming behavior of the cryoinjured fish. Swimming speed was assessed in fish one hour prior to analgesic treatments and cryoinjury (−1H), and at 2, 6, 24, and 48H after cryoinjury in injured untreated control fish (n=9, black circles), in injured lidocaine-treated fish (n=9, red squares), and injured morphine-treated fish (n=6, dark blue squares). (**F**) Effect of the 6H morphine treatment on the swimming behavior of the cryoinjured fish. Swimming speed was assessed in fish prior to morphine treatment and cryoinjury at -1H time point, and at 2, 6H after cryoinjury in injured untreated control fish (n=18, black circles), and in injured morphine-treated fish (n=18, light blue squares). Mean ± SEM. Two-way ANOVA with Sidak’s multiple comparison test (*P*>0.05*). Dashed lines in **E, F** correspond to injury.

In our study, we tested the systemic application of two analgesics, lidocaine (3mg/l) (Schroeder and Sneddon, 2017) and morphine (1.5 mg/l) (Khor et al., 2011), as pain-relieving agents after heart cryoinjury and analyzed the fish swimming behavior as a surrogate to assess the animal welfare. We evaluated the effect of a long-term treatment of 48 hours (H) after the cryoinjury for both analgesics and a short-term treatment of 6H for morphine only (Figure 1C). Our data showed the zebrafish changed their swimming behavior after cryoinjury reflecting the possible noxious effects associated with this procedure. The administration of morphine but not lidocaine alleviated the changes in the swimming activity as a sign of pain relief. Importantly, we also determined the impact of these analgesics on the heart regenerative process. Using histology, scRNA sequencing and RNAscope *in situ* hybridization we demonstrated the short-term morphine treatment does not impede the heart regenerative process. Taken together, we propose to refine the cryoinjury procedure by administering morphine at 1.5 mg/l for a period of 6H after cryoinjury and recommend implementing this practice by researchers in the field. Our data also point out other invasive procedures to be reevaluated to ensure the best practices for zebrafish welfare.

## Results

### Morphine but not lidocaine affects the zebrafish behavior after cryoinjury

Reportedly, the swimming behavior may be used as a surrogate to quantitate fish welfare (Deakin et al., 2019; C. I. M. Martins et al., 2012). We monitored the changes in the swimming speed reflecting potential post-cryoinjury pain for the whole period of the systemic analgesic administration, while the impact of the analgesic treatment on the regenerative process was examined at 7, 15, and 30 days post injury (dpi) (Figure 1C). First, we set out to determine whether lidocaine and morphine have any impact on zebrafish behaviour in a steady state (i.e. without cryoinjury). We monitored the swimming speed for a period of 48H in the untreated control group and compared it to the lidocaine- (3 mg/l) and morphine-treated (1.5 mg/l) group. Our results do not show any significant differences in swimming speed between the untreated control fish, lidocaine- and morphine-treated fish (Figure 1D). Nonetheless, when the test groups are compared individually, the lidocaine-treated fish swim at 6H significantly slower than at 2H (Figure S1A), consistent with its sedative effect. In agreement with the studies showing high variability in locomotor behavior both in larvae (Fitzgerald et al., 2019) and in adults (Lange et al., 2013), we also noticed a high variability in the swimming speed between the individual fish across all groups, highlighting the complexity of behavioral studies.

To determine whether the cryoinjury procedure may cause adverse nociceptive effects, we compared the swimming behavior of fish before and after the cryoinjury, and observed a significant reduction in the swimming speed, suggesting the procedure may inflict distress and/or pain (Figure 1E, F, black filled circles, Figure S1B). We surmise the analgesic treatment should improve the swimming behavior of cryoinjured fish. We therefore quantitated fish swimming speed after the cryoinjury procedure followed by a treatment with either lidocaine or morphine over period of 48H at different time points: 2, 6, 24, and 48 hours post injury (hpi) (Figure 1E). When comparing all three groups together, the observable differences occurred only at 2H of analgesic treatment (i.e. 2 hpi), in the later time points all three groups behaved similarly. While lidocaine treatment did not improve the swimming speed after cryoinjury, there was a noticeable, although not statistically significant, difference in the swimming behavior of the morphine-treated fish (Figure 1E). The data indicates morphine might be a potentially good candidate for the analgesic treatment to reduce pain and distress after cryoinjury in adult zebrafish.

Since the major impact of the analgesic treatment could be observed only in immediate hours after the cryoinjury procedure, we decided to repeat the experiment to test the effect of a short-term 6H morphine treatment after cryoinjury with a higher number of fish per group (Figure 1C, F). Our data show the morphine treatment significantly alleviates the pain-related effects after cryoinjury at 2 hpi and may be beneficial up to 6 hpi (Figure 1F).

Altogether, our data show the cryoinjury may cause adverse effects associated with pain that can be alleviated with the morphine treatment (1.5 mg/l) up to 6 hpi. In contrast, lidocaine treatment (3 mg/l) does not consistently show improved animal welfare and over longer periods its effect appears to be sedative.

### Lidocaine treatment delays the heart regeneration process

Zebrafish heart regeneration is a very dynamic process and can be divided into three overlapping phases (inflammatory, reparative, regenerative), which can be distinguished using histology (Chablais et al., 2011). We focused on the reparative and regenerative phases, characterized by collagen and fibrin deposition at 7 dpi, matrix degradation at 15 dpi and myocardium replacement followed by the completion of the process at 30dpi, respectively (Figure 1C). We assessed injured hearts isolated from the untreated (Figure 2A-A’’), lidocaine-treated for 48H (Fig. 2B-B’’), morphine-treated for 48H (Fig. 2C-C’’), and morphine-treated for 6H (Fig. 2D-D’’) fish, using the histological dye Acid Fuchsin Orange G (AFOG) staining. While the regenerative capacity of hearts from injured untreated and morphine-treated fish is comparable at all tested time points, hearts from the lidocaine-treated fish show a marked delay at every phase of the regenerative process, with a visibly larger injured area (IA) filled with fibrotic and collagen matrix, still present at 30 dpi (Fig. 2A-D). The quantification of the IA at 7 dpi, 15 dpi and 30 dpi in all tested conditions corroborated our initial observation, with the most significant difference between lidocaine-treated and untreated hearts occurring at 7 dpi (Fig. 2E), suggesting the lidocaine treatment may interfere with the reparative phase of the regenerative process. Altogether, in addition to its sedative effects, lidocaine delays the heart regeneration in zebrafish, making it unsuitable as an analgesic to improve the fish welfare after cryoinjury. On the contrary, a 6H morphine treatment post-cryoinjury seems to be sufficient to alleviate the adverse impact of the procedure, while not interfering with heart regeneration. We therefore used the 6H morphine treatment (1.5 mg/l) after cryoinjury in all subsequent experiments and refer to this experimental condition as morphine-treated hearts.

**Figure 2.**
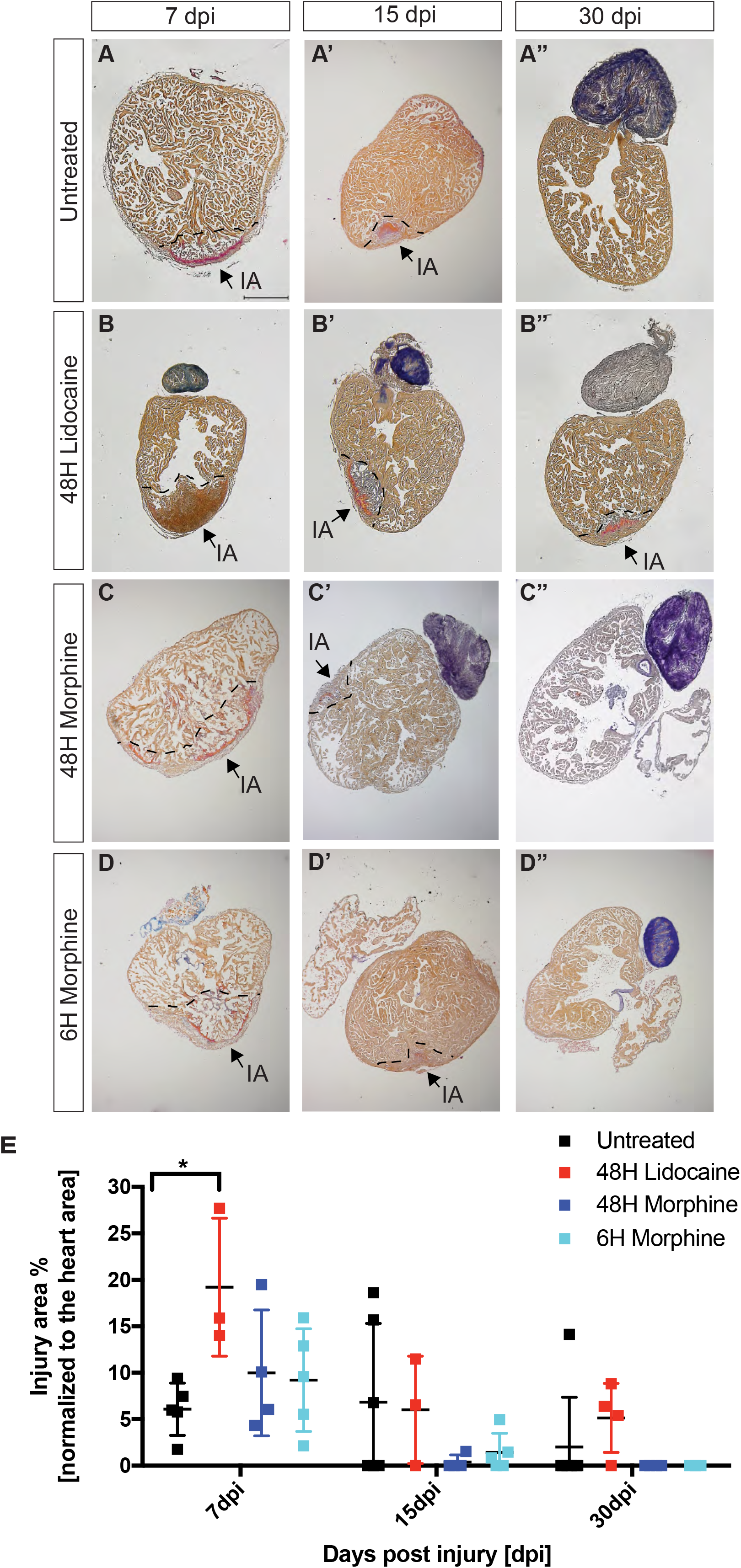
Lidocaine but not morphine treatment delays heart regeneration. (**A-D**) Effect of analgesics on the regenerative capacity was compared at 3, 7, 15, and 30 dpi between untreated injured hearts (**A-A’’**), lidocaine-treated (**B-B’’**) and morphine-treated (**C-C’’**) hearts for a period of 48H, and morphine-treated hearts for a 6H period (**D-D’’**). The histological dye Acid Fuchsin Orange G stains healthy myocardium in orange, fibrin in red, and collagen in blue. IA, injury area. Scale bar, 300 μm. (**E**) Quantification of the injury area in % normalized to the total ventricular area, n(untreated)=5, n(lidocaine_48H_)=3, n(morphine_48H_)=4, n(morphine_6H_)=5. Two-way ANOVA with Sidak’s multiple comparison test (*P*>0.05*).

### Morphine treatment does not affect the heart regeneration process

Our behavioral and histological analysis indicated morphine might be the analgesic of choice to effectively manage noxious effects associated with the cryoinjury procedure in zebrafish while at the same time minimally impacting the regeneration machinery. To gain more insights into any possible effects of the morphine on heart regeneration we deployed single-cell RNA sequencing (scRNAseq) to determine whether morphine treatment induces any changes in the cell type diversity. We single-cell sequenced the transcriptomes of the injured hearts isolated at 3, 7, and 15 dpi from either untreated or morphine-treated fish. We detected all the major cardiac cell types in the clustered cellular transcriptomes (Figure 3A). Comparing the cellular diversity between the transcriptomes of untreated and morphine-treated fish revealed an almost perfect overlap between the two datasets, with neither additional nor missing cell types (Figure 3B). Moreover, direct comparison of the cell types as percentage of total cell count showed all the identified cell types were represented in comparable numbers between untreated and morphine-treated fish (Figure 3C). Plotting the mRNA expression of the individual genes with a defined role in regeneration including *aldh1a2, cxcl12a, postnb, gata4, cxcr4b, fn1a, col1a1a*, and *tgf1b* (Bensimon-Brito et al., 2020; Gupta et al., 2013; Itou et al., 2012; Kikuchi et al., 2011; Kim et al., 2010; Liu et al., 2018; Moyse and Richardson, 2020; Sánchez-Iranzo et al., 2018; Wang et al., 2013) did not reveal any significant differences between the untreated and morphine-treated fish (Figure 3D). Quantifying the relative mRNA expression of a subset of these genes further corroborated our scRNAseq data (Figure 3E). Taken together, our systematic analysis of the single cell transcriptomes revealed a 6H morphine treatment of fish after cryoinjury does not seem to affect the regenerative machinery on the cellular and the molecular level.

**Figure 3.**
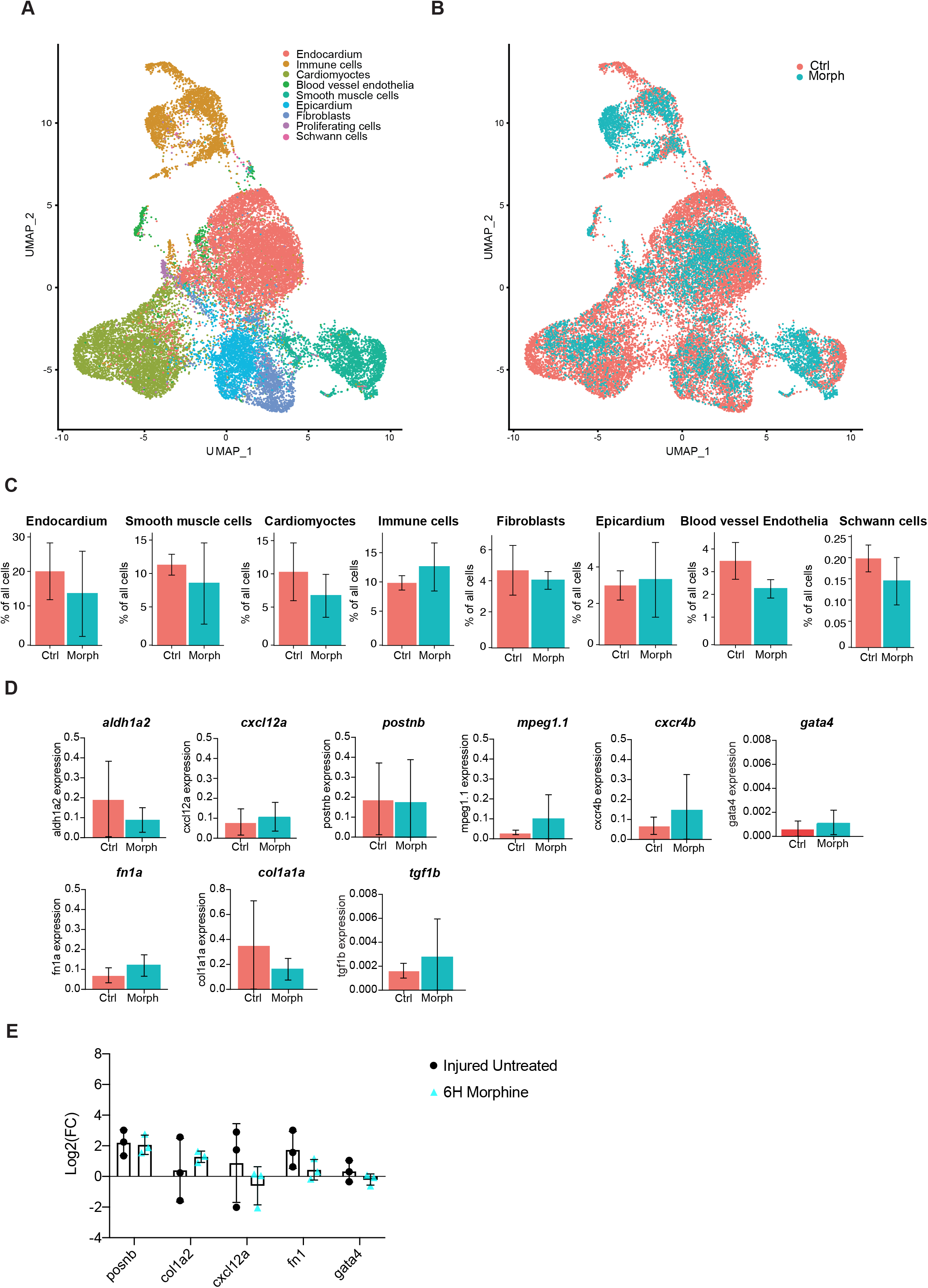
Morphine treatment does not affect gene expression during heart regeneration. (**A, B**) UMAP representation of scRNA sequencing from pooled injured untreated hearts and injured 6H morphine-treated hearts collected at 3, 7, and 15 dpi, n(untreated)=3, n(6H-morphine)=3. All identified cell types (**A**) are present in both untreated and morphine-treated hearts (**B**). (**C**) Comparison of means of individual cell type groups as % of all cells between untreated hearts and 6H morphine-treated hearts. Unpaired student t-test with Welch’s correction (*P*>0.05). *P* values show no significant difference. (**D**) Comparison of mean gene expression of selected genes involved in the regeneration between untreated hearts and 6H morphine-treated hearts. Mean ± SEM. Unpaired student t-test with Welch’s correction (*P*>0.05). *P* values show no significant difference. (**E**) Comparison of relative mRNA expression of genes involved in heart regeneration between untreated and morphine-treated hearts. Data plotted as log_2_ of fold change (FC) ± SD of N=3 experiments. Unpaired student t-test with Welch’s correction (*P*>0.05).

To further probe the regenerative process, we focused on cell proliferation as a paradigm of the zebrafish response to heart injury. The percentage of proliferating cells over total cell count as well as mRNA expression of *pcna*, was similar between untreated and morphine-treated hearts (Figure 4A). To validate this data, we performed RNAscope *in situ* hybridization using *pcna* as a marker to visualize proliferative cells at 7 and 30 dpi (Figure 4B, C). At 7 dpi, IA is characterized by high collagen deposition and concomitant induction of cardiomyocytes proliferation at the injury border zone; these features can be visualized by the *collagen I* and *vmhc* (ventricular myosin heavy chain) expression, respectively (Figure 4B). Both untreated and morphine-treated hearts show the proliferating cardiomyocytes labeled through the co-localization of *vmhc* and *pcna* (Figure 4B, arrows in merge panel). Additionally, the proliferation of other cell types, including fibroblasts (co-localization of *collagen I* and *pcna*), is also readily visible. In agreement with previous reports, the regeneration process was mostly finished at 30 dpi as seen by the absence of the *pcna* marker and low levels of *collagen I* that is confined to the epicardial layer in both untreated and morphine-treated hearts (Figure 4C).

**Figure 4.**
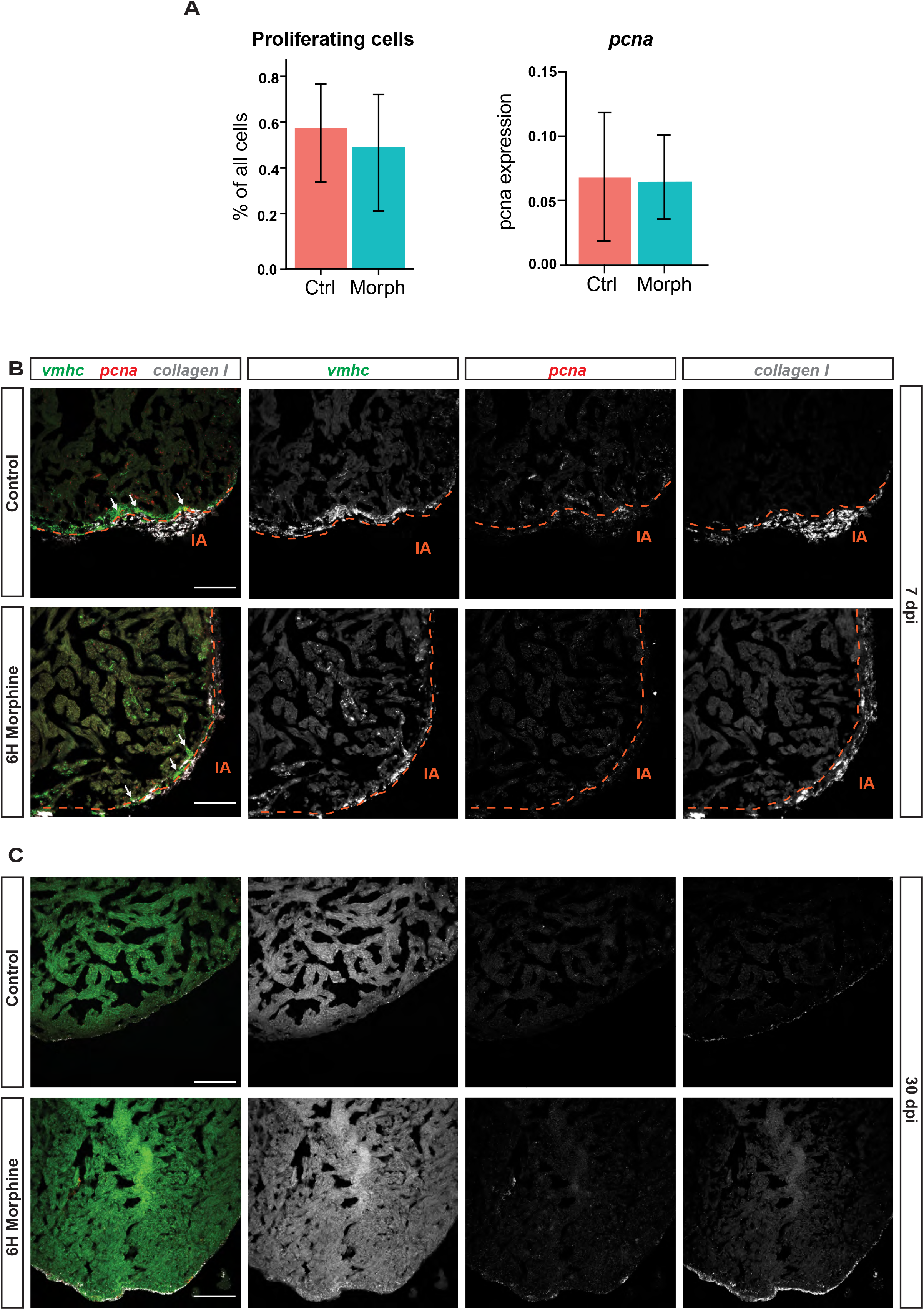
Morphine treatment does not impair cell proliferation during heart regeneration. (**A**) Comparison of means of proliferating cells as % of all cells and mean gene expression of *pcna* between untreated hearts (n=3) and 6H morphine-treated hearts (n=3). Mean ± SEM. Unpaired student t-test with Welch’s correction (*P*>0.05). (**B, C**) Using RNAscope *in situ* hybridization, the amount of proliferating cells labeled with *pcna* RNAscope probe (red) is comparable between untreated hearts and 6H morphine-treated hearts at 7dpi (**B**). Injury area (IA) was detected using *collagen I* (grey), and cardiomyocytes with *ventricular myosin heavy chain, vmhc*, (green) probe. At 30dpi (**C**) there are not any proliferating cells detected in both untreated hearts and 6H morphine-treated hearts. Scale bar, 100 μm. Arrows indicate proliferating cardiomyocytes.

In summary, we show morphine at the concentration of 1.5 mg/l administered for the first 6H post-cryoinjury appears to alleviate the noxious effects associated with the procedure. Moreover, we could not observe any significant impairment in the heart regeneration after this short-term exposure, neither at the cellular nor at the molecular level.

## Discussion

Our study highlights two points: 1) the zebrafish significantly change their swimming behavior after cryoinjury, indicating this procedure affects them adversely, and may cause stress and/or pain; 2) morphine can be administered to reduce post-surgical pain in adult zebrafish without impeding the regenerative process. Our study highlight the great need to systematically study the effects of the analgesics in adult zebrafish, both on their ability to manage stress and/or pain, and on the physiological process studied.

Zebrafish has been used as a model in heart regeneration studies for almost 20 years (González-Rosa et al., 2017). During this time a number of methods to induce heart injury have been developed to best mimic the myocardial infarction in humans (González-Rosa et al., 2017). In this study, we used a widely established protocol in heart regeneration field to induce an ‘ischemia-like’ necrotic and apoptotic cell death across the ventricle of adult fish using a cooled cryoprobe, mimicking human infarction (Chablais et al., 2011; González-Rosa and Mercader, 2012). Although the injury is well tolerated by animals, with a high survival rate of 95%, whether the animals sense the adverse effects associated with stress and/or pain had not been thoroughly investigated.

The question whether fish can sense pain has been rather controversial. Nevertheless, fish respond to noxious stimuli similarly as other animals (Chatigny et al., 2018; Sneddon, 2015), which warrants revision of surgical protocols to manage potential adverse effects associated with such procedures. Studies that systematically address the pain management in zebrafish, and fish in general, have only recently received much needed attention (Chatigny et al., 2018; Deakin et al., 2019; Lopez-Luna et al., 2017; Schroeder and Sneddon, 2017; Sneddon, 2012). In this study, we used the swimming activity as a surrogate to assess the level of stress and/or pain after the cryoinjury by adapting previously reported assays (Deakin et al., 2019). Our data show the significant decrease in the swimming activity after cryoinjury at least up to 6 hpi, indicating this procedure could be considered as a potential noxious stimulus and requires revising, assuring any possible pain and discomfort of animals can be reduced to minimum.

Currently, there are no measures to minimize pain after heart injury procedures in adult zebrafish. In this study, we have tested two analgesics, lidocaine and morphine and administered them systemically by immersion. We have opted for assessing only one concentration per analgesic based on previous reports (Deakin et al., 2019; Khor et al., 2011; Lopez-Luna et al., 2017; Schroeder and Sneddon, 2017), but tested different durations of treatments. The usage of lidocaine as a systemic pain reliever is rather popular (Mao and Chen, 2000). However, our data indicate lidocaine in zebrafish causes more sedative-like effects shown by the tendency of lidocaine-treated fish to swim slower. Systemic administration of morphine in fish has been associated with beneficial effects (Chatigny et al., 2018). Indeed, we have determined morphine, but not lidocaine improves the swimming activity of zebrafish after cryoinjury; this effect is statistically significant up to 2 hpi and persists to improve the swimming activity up to 6 hpi.

The choice of an appropriate analgesic should be always considered pertaining the physiological processes being studied. Since the administration of both lidocaine and morphine has been associated with undesired side effects (Chatigny et al., 2018), we have not only assessed their effect on the fish swimming behaviour after the cryoinjury, but have also determined their impact on the regenerative process. Our histological analysis demonstrated lidocaine, but not morphine reduces the capacity of the zebrafish heart to regenerate. As lidocaine inactivates the fast voltage-gated Na^+^channels, one possible explanation could be a defective wound healing process, which partially relies on the restoration of transepithelial electrical gradients (Dubé et al., 2010; Zhao et al., 2006). Importantly, our detailed comparison of the cellular and molecular responses between control and morphine-treated fish determined major differences neither in the cellular composition nor at the transcriptional level. We have tested not only genes with established roles in the heart regeneration (Bensimon-Brito et al., 2020; Gupta et al., 2013; Itou et al., 2012; Kikuchi et al., 2011; Kim et al., 2010; Liu et al., 2018; Moyse and Richardson, 2020; Sánchez-Iranzo et al., 2018; Wang et al., 2013), but have also probed the proliferative response of CM, which is a hallmark of the zebrafish heart regeneration. We, however, advise caution as the earliest assessed transcriptomes were of hearts isolated at 3dpi, and thus we cannot rule out morphine effects on the early wound healing and on the immunological response.

Taken together, morphine should be considered as an analgesic of choice to reduce stress and/or pain after cryoinjury in adult zebrafish. We recommend the systemic administration of morphine at 1.5 mg/l for 6 hours as a method refinement. The cryoinjury procedure under these conditions, i.e. with proper anesthesia followed by analgesia, should be considered as a moderate burden for the animal.

## Methods

### LEAD CONTACT AND MATERIALS AVAILABILITY

Further information and requests for resources and reagents should be directed to and will be fulfilled by the Lead Contact, Daniela Panáková. All reagents generated in this study are available from the Lead Contact without restriction.

## KEY RESOURCES TABLE

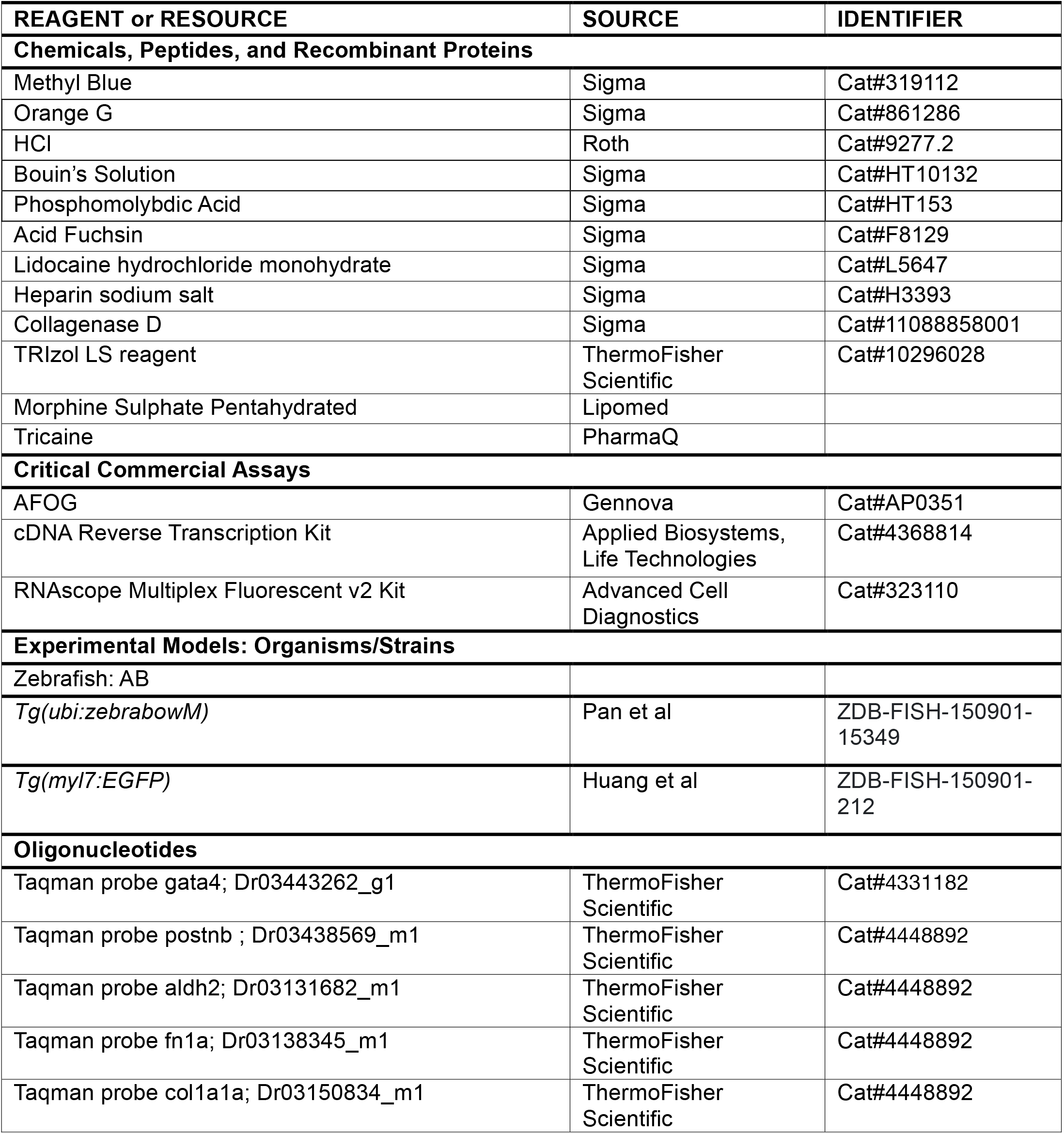
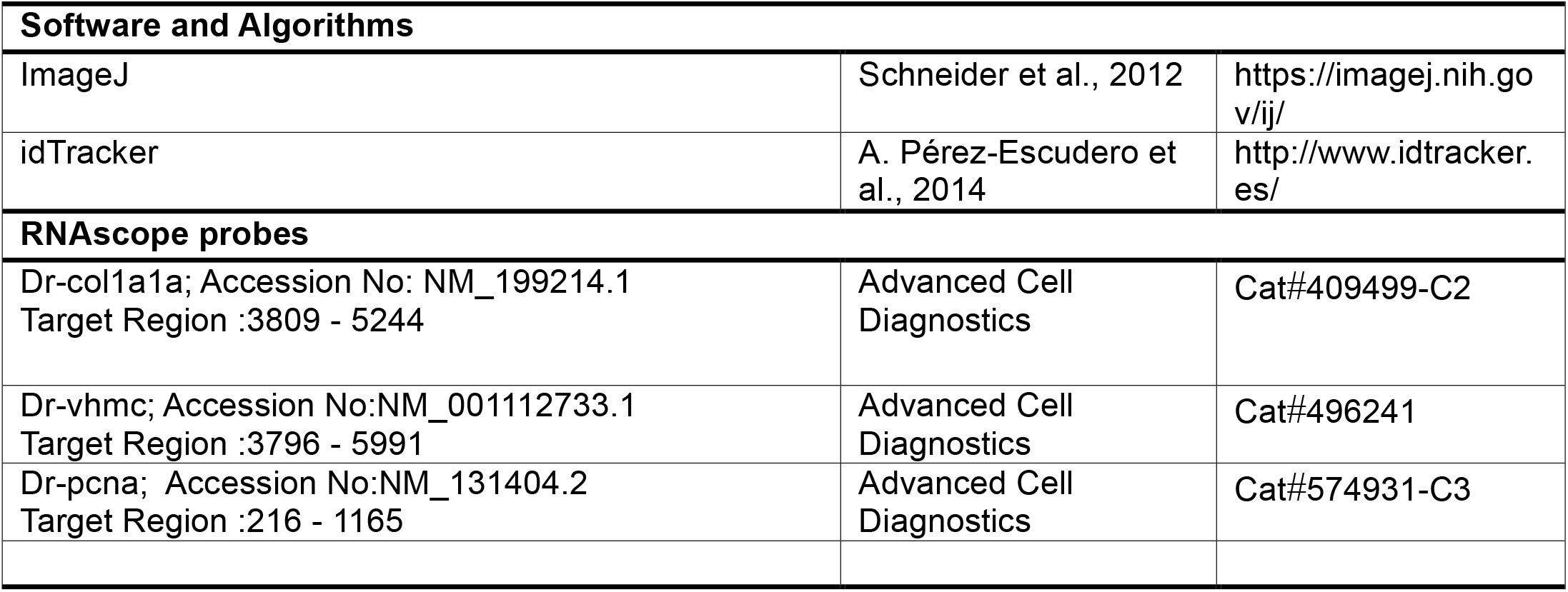

## EXPERIMENTAL MODEL AND SUBJECT DETAILS

### Animal Studies

Zebrafish were bred, raised, and maintained in accordance with the guidelines of the Max-Delbrück Center for Molecular Medicine and the local authority for animal protection (Landesamt für Gesundheit und Soziales, Berlin, Germany) for the use of laboratory animals based on the current version of German law on the protection of Animals and EU directive 2010/63/EU on the protection of animals used for scientific purposes. In addition, housing and breeding standards followed also the international ‘Principles of Laboratory Animal Care’ (NIH publication no. 86-23, revised 1985). Zebrafish strain *AB* and transgenic lines used in this study included *Tg*(*myl7:EGFP*)^twu34^ (Huang et al., 2003) a*nd Tg(ubi:zebrabow-M)*^*a131Tg*^ (Pan et al., 2013). Embryos were kept in E3 embryo medium (5 mM NaCl, 0.17 mM KCl, 0.33 mM CaCl_2_, 0.33 mM MgSO_4_, pH 7.4) under standard laboratory conditions at 28.5 °C. Adult zebrafish of both genders, aged between 6 months and a year, and bigger than 3 cm were used.

### Drug Treatments

Fish were treated with either 3 mg/l of lidocaine hydrochloride monohydrate (Sigma) or 1.5mg/l morphine sulphate pentahydrated (Lipomed) dissolved in system water. The concentration of the analgesics was based on the previously published reports (Khor et al., 2011; Schroeder and Sneddon, 2017). Fish were held in individual tanks for the duration of treatment, between 2 to 48 hpi, with exchange of the drug dissolved in fresh system water twice a day.

### Analysis of zebrafish swimming activity

Fish were monitored by recording their swimming activity with a video camera positioned over the tanks for a 3-minute period at the following time points: 1 hour (H) before the treatment, 2, 6, 24 and 48 H after the treatment. The swimming speed (mm/sec) was scored using the software idTracker, calculating the change in position in two dimensions (Correia et al., 2011; Pérez-Escudero et al., 2014).

### Cryoinjury procedure

Cryoinjury was performed as previously described (Chablais et al., 2011; Chablais and Jazwinska, 2012; González-Rosa and Mercader, 2012). Briefly, fish were pre-sedated in water containing 0.03 mg/ml Tricaine (PHARMAQ, pH 7). Concentration was then increased to 0.168 mg/ml for anesthesia. Fish were placed with the ventral side up to access the heart; a small incision was made through the body wall and the pericardium using microdissection forceps and scissors. Once the pericardial sac was opened, the heart ventricle was exposed by gently compressing the abdomen. Excess water was carefully removed by blotting with tissue paper, not allowing fish skin to dry. Then, a stainless steel cryoprobe precooled in liquid nitrogen was applied to the ventricular wall for 20 seconds. Fish were then placed in a tank of fresh system water with or without analgesic; reanimation was enhanced by the gills oxygenation where water around gills was aerated by pipetting for a couple of minutes.

### Histological staining, analysis and imaging

For the analysis of the injury areas, animals were humanely killed at different times post-injury by placing them in ice cold water of 0-4°C for 20 minutes. Hearts were dissected and incubated in 2U/ml heparin and 0.1KCl in PBS for 30min. Cryosamples were fixed in 4% PFA in PBS overnight at 4°C, washed in PBS for 3 x 10 min, and incubated overnight in 30% sucrose in PBS. Samples were then frozen in Tissue-Tek O.C.T Compound (Sakura) on the dry ice. Tissue was cut at 7µm on a cryostat (Leica) using Superfrost slides (ThermoFisher Scientific). Connective tissue was stained using acid fuchsin orange G (AFOG). In brief, slides were dried for 30min at RT. Next, slides were incubated at Bouin Solution (Sigma-Aldrich) for 2H at 60°C and left for overnight incubation under the hood. Slides were washed for 30min under running water and incubated for 7min in 1% phosphomolybdic acid (Sigma-Aldrich). Samples were washed for 3min in running ddH_2_O and incubated with AFOG solution (self-made, Sigma-Aldrich) for 3min. Slides were washed until clear with running ddH_2_O and rehydrated with 70%, 94%, 2x 100% ethanol and 2x 5min xylol (Sigma-Aldrich). Slides were mounted with xylene mounting medium (Merck) and let dry overnight under the hood. For the analysis of injury size, the total ventricular tissue area and injury area (IA) on multiple slides were measured. Imaging was done using a Keyence Microscope BZX800 and analyzed with ImageJ/Fiji.

### Preparation of single-cell suspensions

Adult zebrafish were humanely killed at different times post-injury by placing them in ice cold water of 0-4°C for 20 minutes. The heart was dissected from the fish and transferred into cold HBSS. A needle and a syringe filled with cold HBSS were used to pierce into the lumen of the heart to thoroughly wash away most of the erythrocytes in the tissue. Afterwards, the tissue was opened carefully with forceps, and the heart tissue was incubated at 37°C for 30 min in 500µl HBSS containing Liberase enzyme mix (Sigma-Aldrich, 0.26 U/mL final concentration) and Pluronic F-68 (Thermo Fisher Scientific, 0.1%), while shaking at 750 r.p.m. with intermittent pipette mixing. After most of the tissue was dissociated, the reaction was stopped by adding 500µl cold HBSS supplemented with 1% BSA. The suspension was centrifuged at 250*g* at 4°C and washed two times with 500µl cold HBSS containing 0.05% BSA, then filtered through a cell strainer of 35µm diameter. The quality of the single cell suspension was then confirmed under the microscope, and cells were counted prior to scRNA-seq library preparation.

### Single cell RNA-seq

Single cells were captured using the Chromium Single Cell 3’ kit (10X Genomics, PN-1000075), according to the manufacturer’s recommendations. We aimed for 10,000 cells per library whenever possible. Both v2- and v3-chemistry were used for data presented here. Samples (3 untreated and 3 morphine-treated hearts of injured fish at 3, 7, 15dpi) were sequenced on Illumina NextSeq 500 2x 75 bp and Illumina HiSeq 2500 2x 100 bp after successful quality control by Bioanalyzer (DNA HS kit, Agilent).

### Mapping and clustering of single-cell mRNA data

A zebrafish transcriptome was created with Cell Ranger 3.0.2. from GRCz10, release 90. Alignment and transcript counting of libraries was done using Cell Ranger 3.0.2. Library statistics are summarized in Table S1. The transcriptome data was filtered, clustered, and visualized using Seurat 3.0 (Stuart et al., 2019).

### Data availability

Sequencing data are deposited on Gene Expression Omnibus, accession number.

### Quantitative Real-Time PCR

Zebrafish hearts were isolated, cut into smaller pieces under the microscope and incubated for 30min in heparin solution (Sigma) followed by incubation for 2 h at 37°C in 0.25 ml collagenase IV (Sigma). 3 hearts per biological replicate were used. RNA from zebrafish hearts was extracted using 0.75 mL TRIzol LS reagent (Invitrogen, Thermo Fisher). RNA was transcribed to cDNA using the High-Capacity cDNA Reverse Transcription Kit (Applied Biosystems, Life Technologies). qRT-PCR was performed using TaqMan probes and solutions (Applied Biosystems). Primer information is available in Key Resource Table.

### RNAscope Multiplex Fluorescent V2 - Immunofluorescence *in situ* hybridization method and imaging

The technique was performed according to the manufacturer’s instructions for fixed frozen tissues (323100-USM). In brief, 7μm cryo sections were dried for 30min at RT and washed for 5min in PBS. Slides were then pretreated with hydrogen peroxide (ACD) for 10min at RT followed by 2x washes in ddH2O. Next, slides were incubated at 99°C for 15min with an antigen retrieval solution (ACD) using a steamer (WMH). Slides were dipped in ddH2O and incubated for 3min in 100% ethanol. After drying, the barrier was drawn and protease III (ACD) was applied for 30min at 40°C in the hybridization oven. Slides were washed 2x in ddH2O and 150 μl of probes (ACD) combinations were applied for hybridization at 40°C for 2 h. Following probes were used (see Key Resource Table). Next, slides were washed 2x 2min in the washing buffer (ACD) and signal amplification using AMP1 (30min), AMP2 (30min) and AMP3 (15min) (ACD) with 2x 2min washes in between was carried out. Signal development was done using TSA plus fluorophores (PerkinElmer) fluorescein, Cyanine 3 and Cyanine 5 in 1:1000 dilution. First, HRP-C1 solution was added for 30min at 40°C, washed 2x 2min in the washing buffer, next chosen fluorophore for 30min at 40°C and in the end blocker for 15min at 40°C were added. The same steps were applied to all three fluorophores. Slides were mounted with ProLong Gold with Dapi Antifade Mountant (ThermoFisher Scientific) and incubated overnight in the dark at RT. Afterwards slides were stored at 4°C. Imaging was done using Zeiss LSM880 confocal microscopy and analysis was performed using ImageJ/Fiji and Photoshop softwares.

## QUANTIFICATION AND STATISTICAL ANALYSIS

Sample sizes are indicated in each figure legend. For the measurement of the injury areas at least 3 biological replicates were used for each condition and each sample has been measured using multiple sections and the average has been calculated. Statistical differences of qPCR expression data were analyzed by unpaired student t-test with Welch’s correction and were considered significant at p<0.05. Injury areas were analyzed by the two-way ANOVA with Sidak’s multiple comparison and considered significant at p<0.05. Swimming speed was analyzed by two-way ANOVA with Sidak’s multiple comparison test. scRNA-seq expression data were analyzed by unpaired student t-test with Welch’s correction. *P* values are indicated in the figure legends and in Table S2.

## Author contribution

Conceptualization, S.L., M.G.S. F.F. and D.P.; Methodology and experimentation, S.L., M.G.S., B.H., A.M.A.A., A.M.M., M.C.; Data analysis, S.L., M.G.S., B.H.; Visualization, S.L., B.H. and D.P.; Writing – Original Draft, S.L. and D.P. with input from all authors; Funding Acquisition, J.P.J. and D.P.; Supervision, J.P.J. and D.P.

## Acknowledgements

We thank Nadia Mercader and Ines J. Marques for training and support, and the Aquatic Facility, Advanced Light Microscopy Facility, and Genomics Platform teams at MDC. Work for this project was supported by German Center for Cardiovascular Research (DZHK, Berlin partner site, project 81Z0100103).

## Figure Legends

**Supplementary Figure 1.**
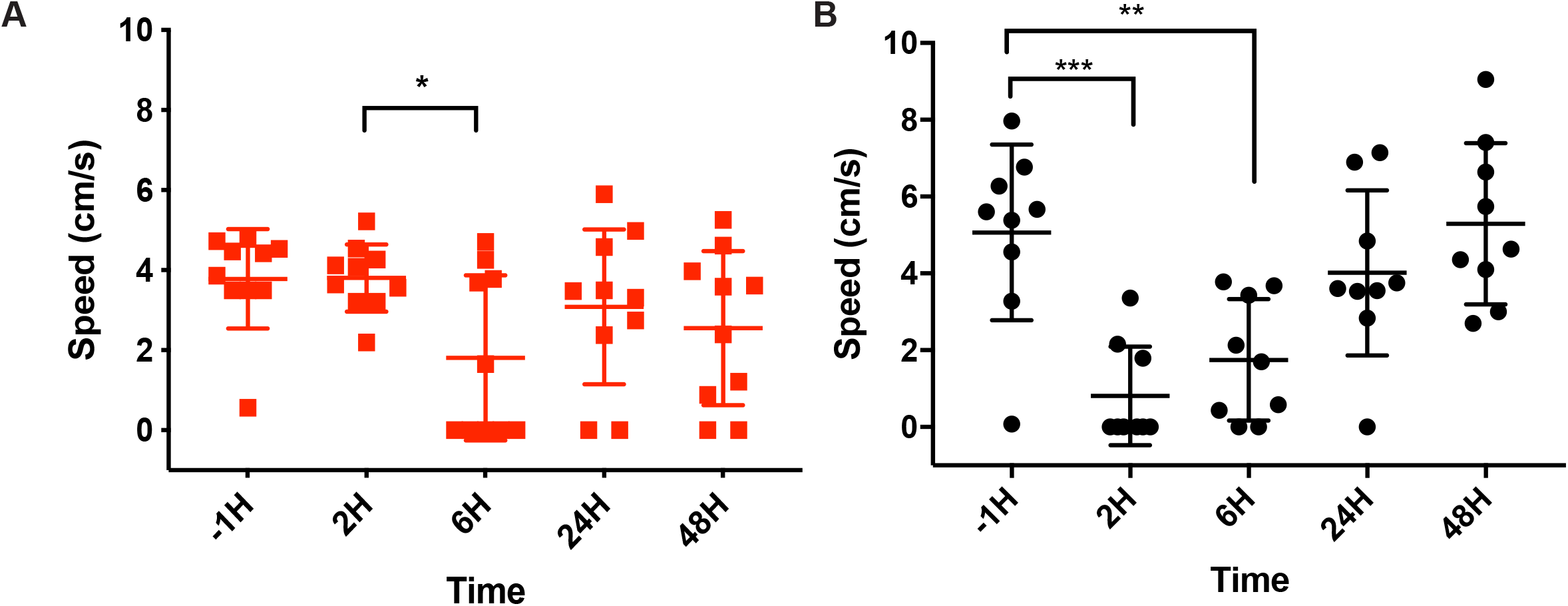
(**A**) Effect of lidocaine treatment on the swimming behavior of the uninjured fish. Swimming speed was assessed in fish one hour prior to analgesic treatments (−1H), and at 2, 6, 24, and 48H (n=10). Mean ± SEM. One-way ANOVA with Bonfferoni’s multiple comparison test (*P* >0.05*; *P* >0.01**) (**B**) Effect of the cryoinjury procedure on the swimming behavior of fish. Swimming speed was assessed in fish one hour prior to cryoinjury (−1H), and at 2, 6, 24, and 48H after cryoinjury (n=9) Mean ± SEM. One-way ANOVA with Tukey’s multiple comparison test (*P*>0.05*).

**Table S1:** scRNA sequencing library statistics

|  | Library name | Days post injury | Average nUMI per cell | Average nGene per cell | Number of cells | 10x Chemistry | Morphine-treated |
| --- | --- | --- | --- | --- | --- | --- | --- |
| 1 | Hr8 | 7 | 2394 | 830 | 5774 | V2 | no |
| 2 | Hr11 | 3 | 1019 | 347 | 10252 | V2 | no |
| 3 | Hr15 | 7 | 1319 | 432 | 5585 | V2 | yes |
| 4 | Hr16 | 15 | 3801 | 832 | 545 | V2 | no |
| 5 | Hr18 | 15 | 2563 | 611 | 643 | V2 | yes |
| 6 | Hr22 | 3 | 6364 | 1279 | 2244 | V3 | yes |

**Table S2:**
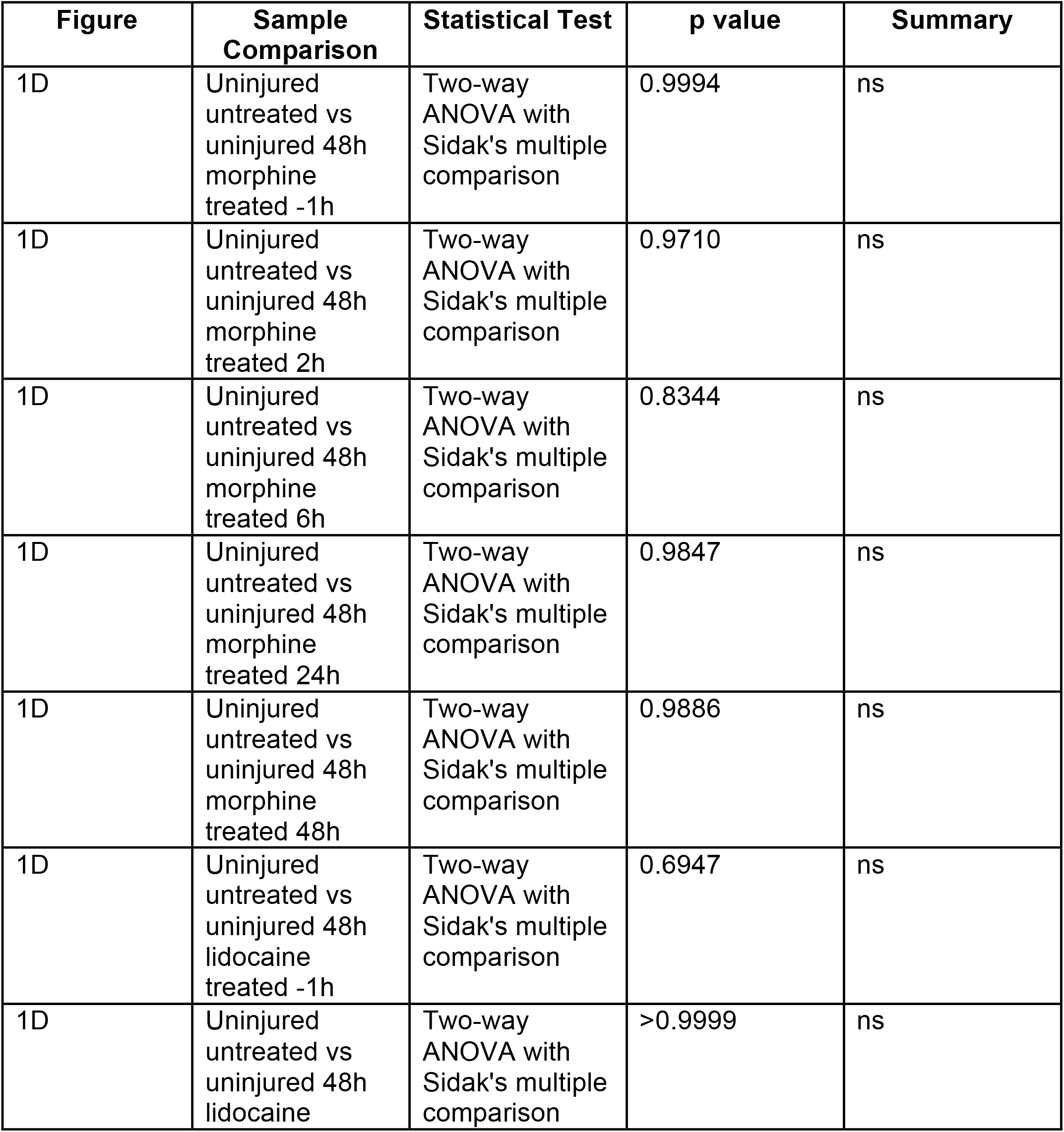
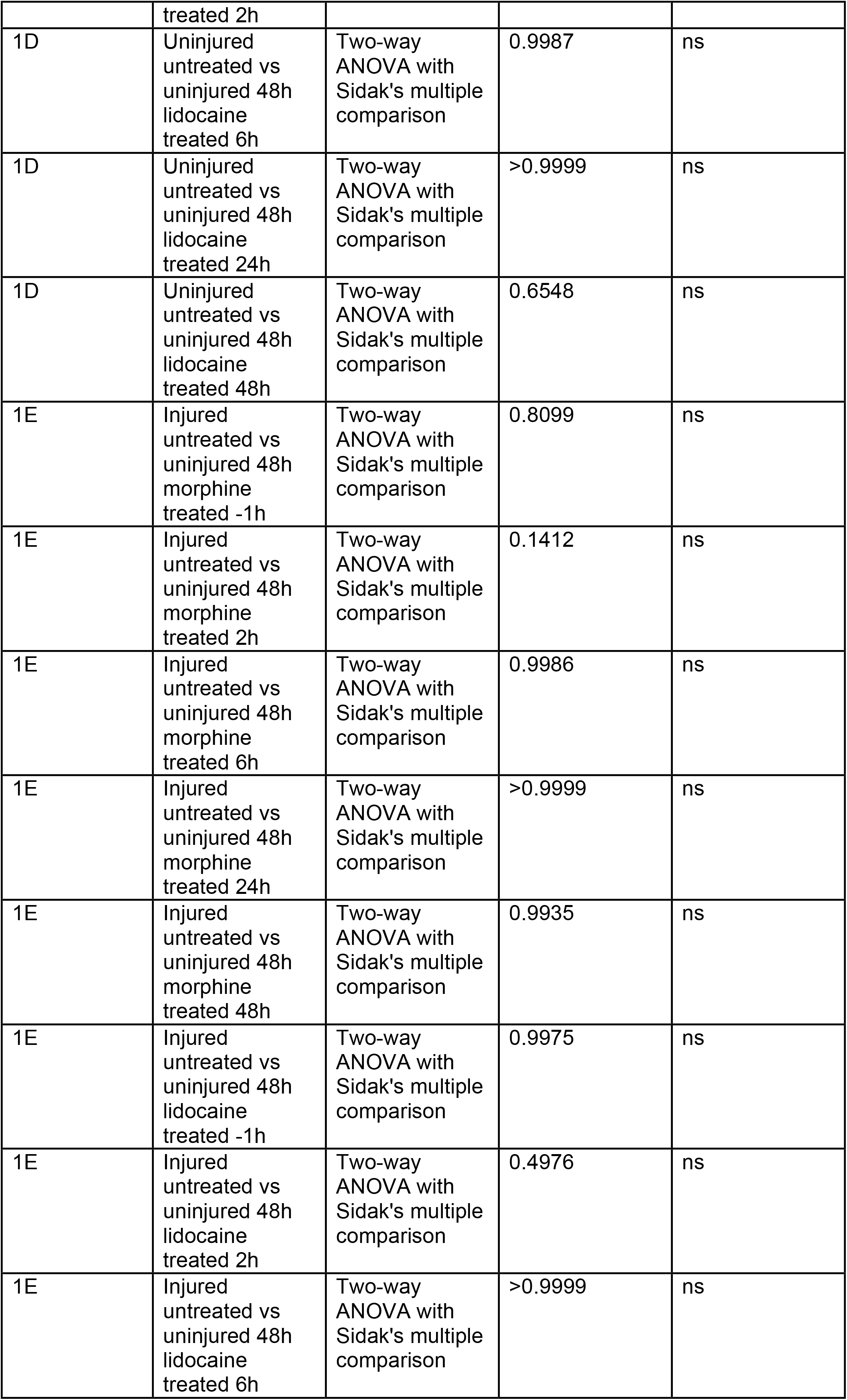
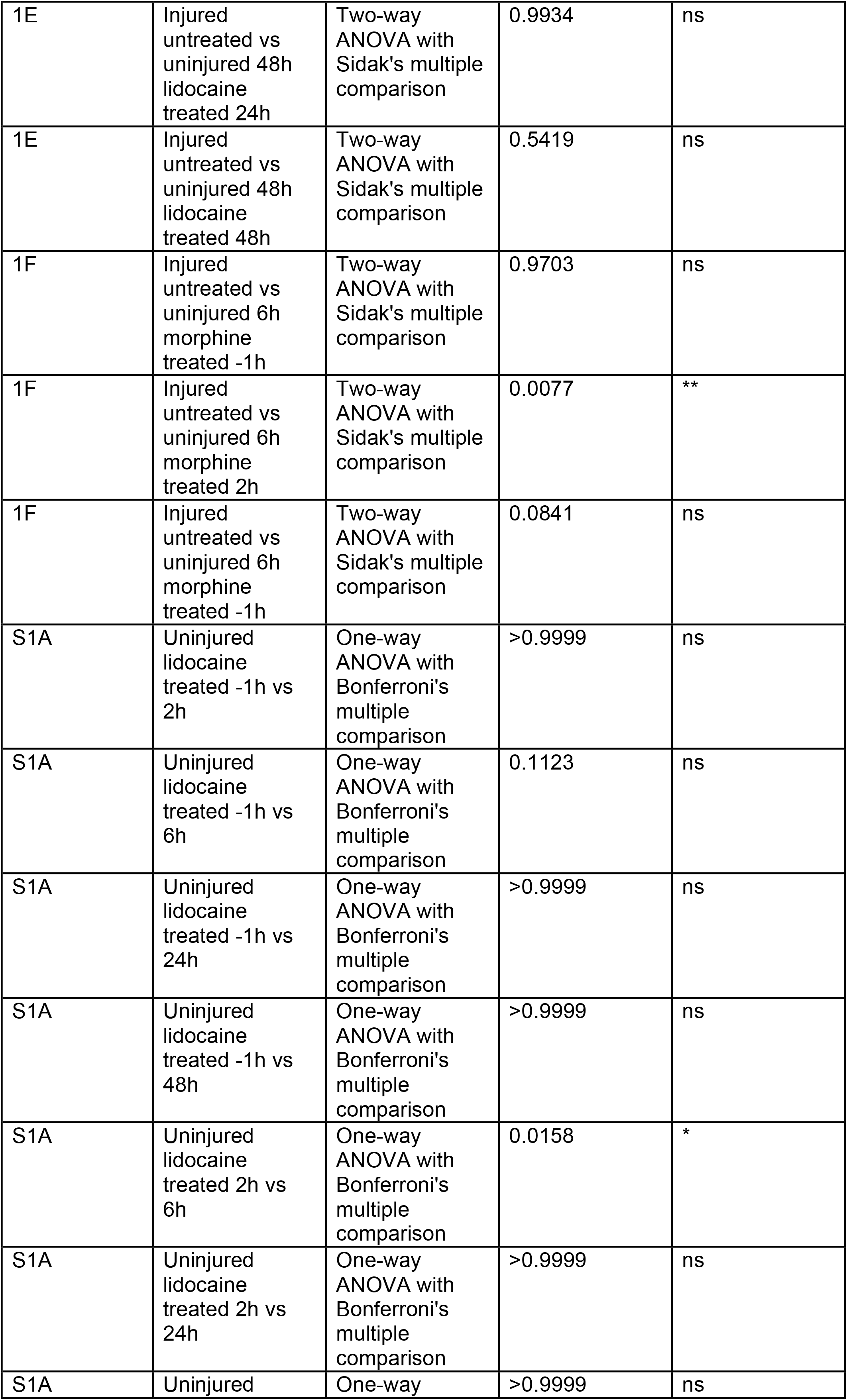
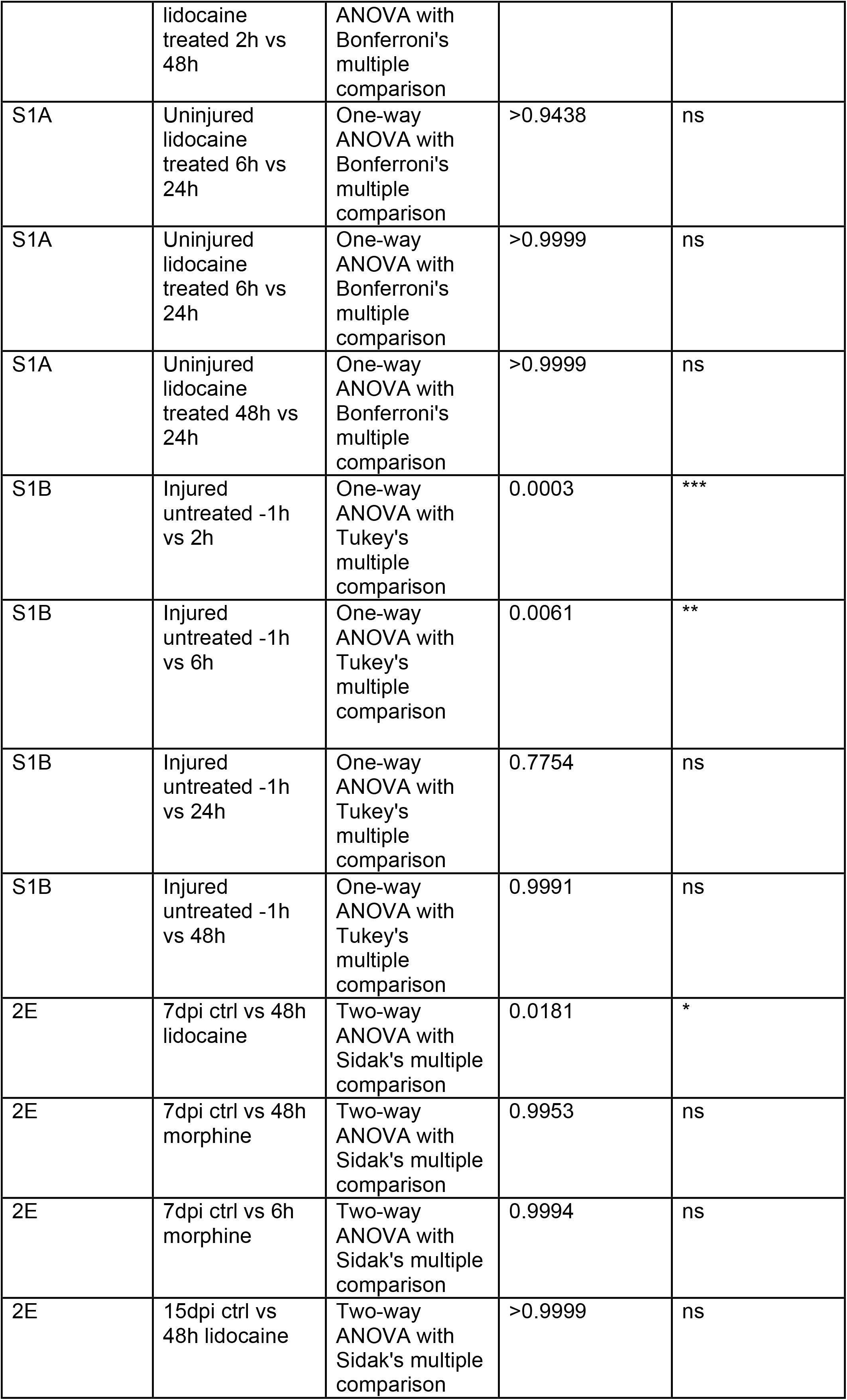
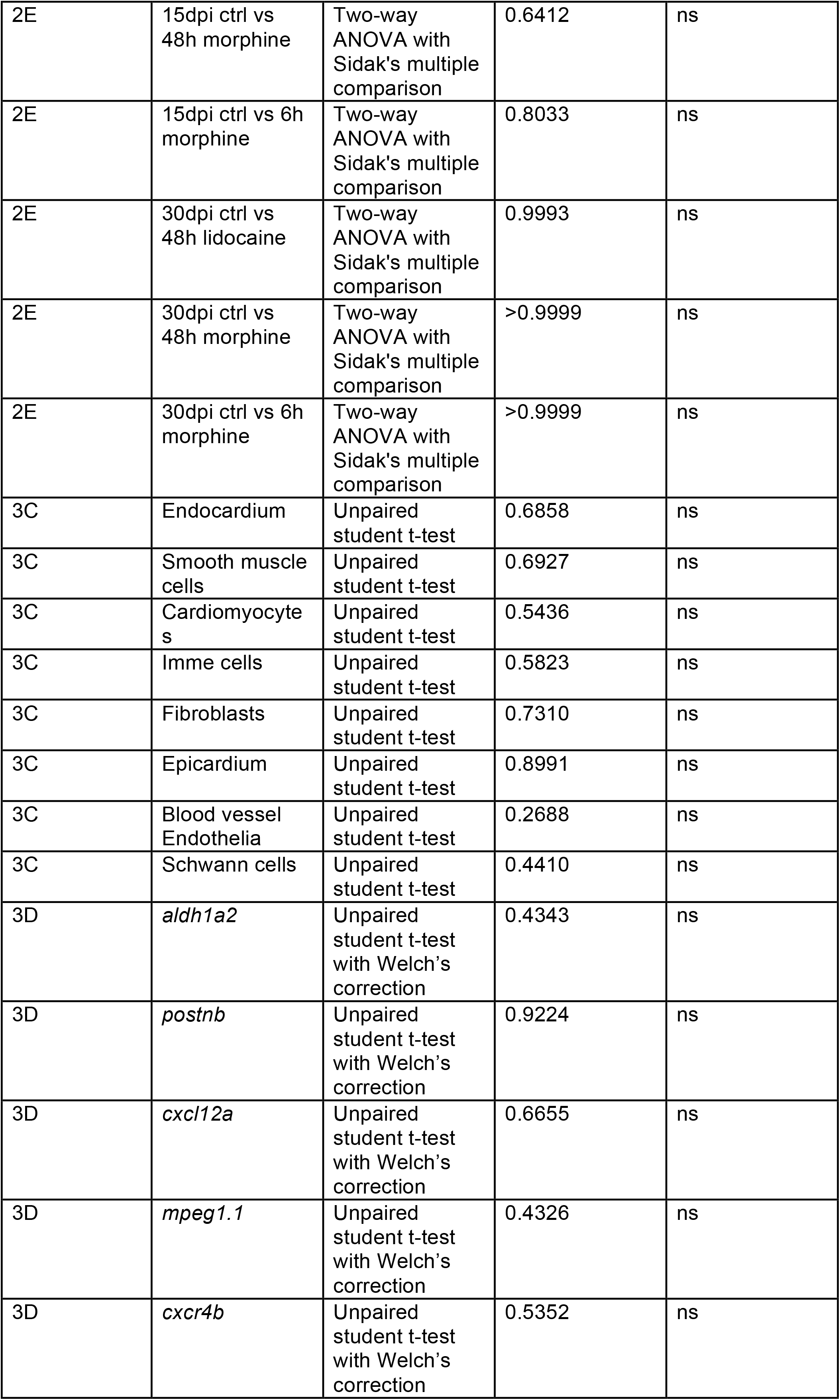
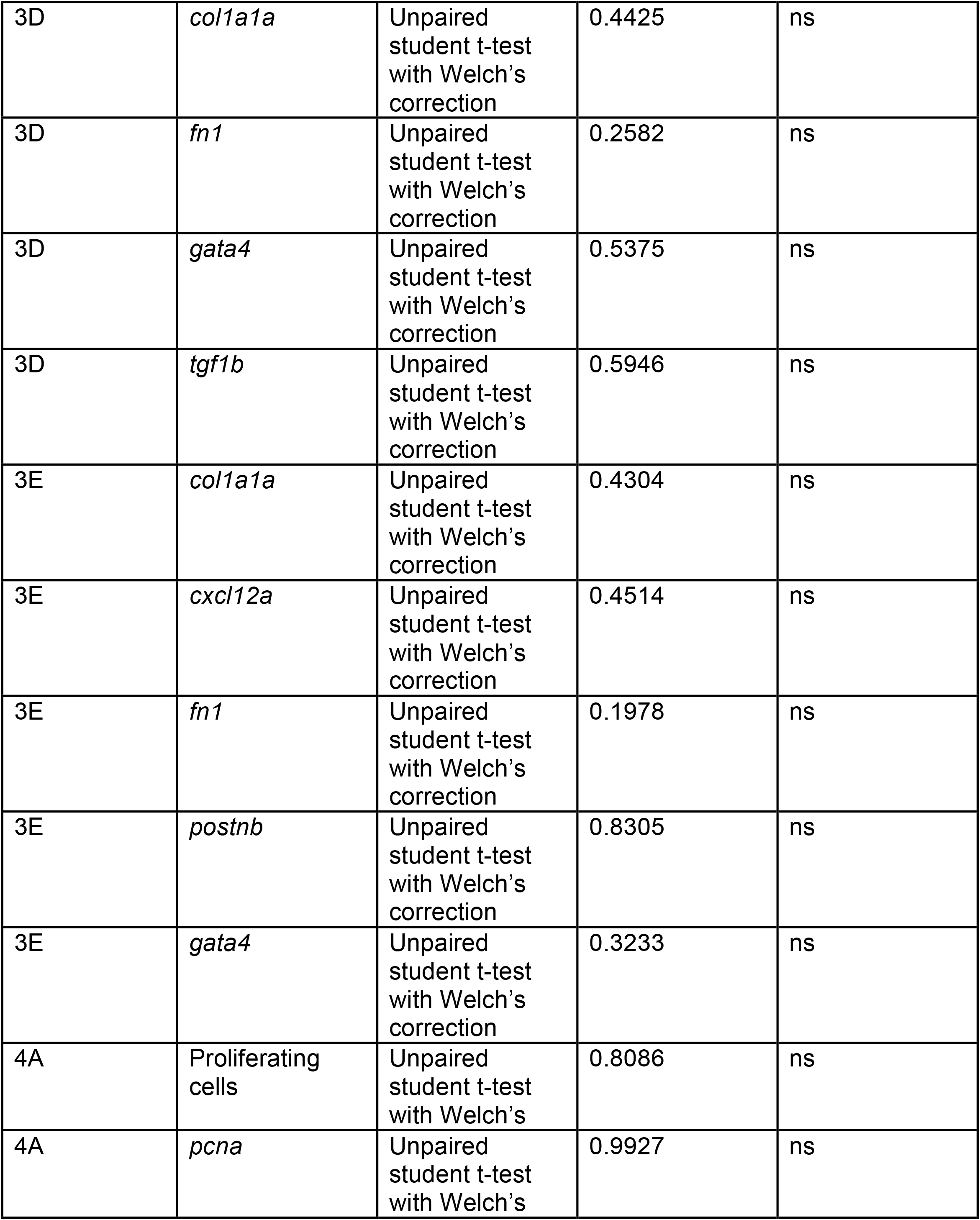
Statistical significance, exact *P* values

## Notes

### Competing Interest Statement

The authors have declared no competing interest.

